# Immuno-informatics Study Identifies Conserved T Cell Epitopes in Non-structural Proteins of Bluetongue Virus Serotypes: Formulation of Computationally Optimized Next-Generation Broad-spectrum Multiepitope Vaccine

**DOI:** 10.1101/2023.11.23.566885

**Authors:** Harish Babu Kolla, Mansi Dutt, Anuj Kumar, Roopa Hebbandi Nanjunadappa, Tobias Karakach, Karam Pal Singh, David Kelvin, Peter Paul Clement Mertens, Channakeshava Sokke Umeshappa

## Abstract

Bluetongue (BT) is a significant arboviral disease affecting sheep, cattle, goats, and wild ruminants, posing serious economic challenges to livestock industry. Control efforts have been hampered by the existence of over 32 distinct BT virus (BTV) serotypes and the absence of broad-spectrum vaccines. Some key non-structural proteins of BTV, including NS1, NS2, and NS3, exhibit notable amino acid sequence conservation. Our findings reveal that mouse MHC class I (MHC-I) CD8+ T cell epitopes are highly conserved in NS1 and NS3, while MHC-II epitopes are prevalent in all the three non-structural NS 1-3 proteins. Similarly, both class I and II Bovine Leukocyte antigen-restricted CD8+ and CD4+ T cell epitopes are conserved within NS1, NS2, and NS3 proteins. To construct *in silico* broad-spectrum vaccine, we subsequently screened these conserved epitopes based on antigenicity, allergenicity, toxicity, and solubility. Modeling and Refinement of the 3D structure models of vaccine constructs were achieved using protein modeling web servers. Our analysis revealed promising epitopes that exhibit strong binding affinities with low energies against two TLR receptors (TLR3 and TLR4). To ensure atomic-level stability, we evaluated the docking complexes of epitopes and receptors through all-atom molecular dynamics simulations (MDS). Encouragingly, our 100 nanoseconds MDS showed stable complexes with minimal RMSF values. Our study offers valuable insights into these conserved T cell epitopes as promising candidates for a broad-spectrum BT vaccine. We therefore encourage for their evaluation in animal models and natural hosts to assess their immunogenicity, safety, and efficacy for field use in the livestock.

## INTRODUCTION

Bluetongue (BT) is a severe arboviral disease primarily afflicting ruminants, especially sheep, and is caused by the Bluetongue virus (BTV), a member of the *Orbivirus* genus within the Reoviridae family (Mertens, Diprose et al. 2004, Matthijnssens, Attoui et al. 2022). BTV’s genome consists of ten segments of double-stranded RNA encoding various structural and non-structural proteins. The structural proteins, organized into core, sub-core, and outer capsid layers, include Viral proteins (VP) 1 through VP7. In contrast, non-structural proteins NS1, NS2, and NS3/3A are positioned alongside the structural components.

Transmission of BT occurs through the bites of *Culicoides* midges, which are blood-feeding insects. Once the virus enters the host, dendritic cells (DCs) recognize, uptake, and migrate to draining lymph nodes (RLNs) (Hemati, Contreras et al. 2009, Umeshappa, Singh et al. 2010, Umeshappa, Singh et al. 2011). In RLNs, the virus replicates before spreading to the spleen, lungs, muscles, and pulmonary artery, leading to severe tissue damage, resulting in edema, vascular thrombosis, hemorrhage, and tissue infarction (Hemati, Contreras et al. 2009, Umeshappa, Singh et al. 2010, Umeshappa, Singh et al. 2011). This devastating infection in animals results in lameness, decreased production, mortality, and significant economic loss (MacLachlan 1994).

Given its high transmission rate, severity, and economic impact, controlling BT spread among ruminants is crucial. However, the virus has evolved into more than 32 strains, rendering conventional serotype-specific vaccines less effective (Rojas 2021). Lessons from influenza vaccine development, targeting conserved immunodominant sequences to combat diverse serotypes, have been informative (Xie 2019, De Jong, Aartse et al. 2020, Lo, Misplon et al. 2021). CIS, being less polymorphic (degree of variation between different serotypes of BTV) and highly conserved among serotypes, can induce cross-protection against all the BTV serotypes. Hence, selecting the ideal antigenic target for identifying CIS and developing a broad-spectrum vaccine is pivotal.

Here, we chose NS proteins as prime vaccine targets due to their high amino acid sequence identity. Employing various bioinformatics approaches, we identified conserved T cell epitopes within NS 1-3 proteins of BTV, highly conserved across 24 serotypes obtained from the NCBI resource. We identified several epitopes that are highly antigenic, non-allergenic, non-toxic, and inducers of IFN-gamma (IFN γ ) for designing a multi-epitope broad-spectrum BTV vaccine. Furthermore, we conducted an in-depth investigation into the vaccine’s capacity to trigger an immune response through molecular docking and molecular dynamics simulations. Our study serves as a proof of concept for future research endeavors aimed at developing effective pan-BTV vaccines capable of conferring cross-protection against all existing BTV serotypes with potential to curb the spread of these virulent serotypes.

## METHODOLOGY

### Sequence retrieval and multiple sequence alignment

The complete amino acid sequences of BTV nonstructural proteins NS1, NS2 and NS3 were retrieved from the available genomes of 24 serotypes of BTV at the National Center for Biotechnology Information (NCBI) taxonomy browser (https://www.ncbi.nlm.nih.gov/Taxonomy/Browser/wwwtax.cgi?id=40051) by using “Bluetongue” as a search keyword. The retrieved amino acid sequences were aligned to identify the conserved CD8+ and CD4+ T cell epitopes. Multiple sequence alignment was performed with ClustalW module in Molecular Evolutionary Genetic Analysis (MEGA-X) software using BLOSUM 65 matrix. The output of alignment files was saved in MEGA format with an extension “.meg” or “.mega”. The amino acid sequence conservation was determined in the MEGA format files of NS1, NS2 and NS3 proteins. The amino acid sequences of these proteins were subjected for BLASTp search against the mouse and bovine proteomes to identify the amino acid similarity of BTV proteins with the host proteome.

### Murine MHC I- and II-restricted epitope prediction

The CD8+ T cell epitopes were predicted for MHC class I alleles in C57BL/6 mice, H2-K^b^ and H2-D^b^. MHC-I-binding epitopes were predicted in NS1, NS2 and NS3 proteins of BTV1 serotype genome with “NetMHCpan BA 4.1” module of the Immune epitope database (IEDB) (http://tools.iedb.org/mhci/) (Vita, Mahajan et al. 2019). Similarly, CD4+ T cell epitopes were predicted corresponding to MHC II H2-IA^b^ allele with “MHCIIpan 4.0 BA” of IEDB tool (http://tools.iedb.org/mhcii/) (Vita, Mahajan et al. 2019). The Half-maximal inhibitory concentration (IC_50_) values were selected as threshold parameters for the selection of epitopes. The immunoinformatics tools use IC_50_ value as a criterion for the epitope prediction which determines the interaction between the epitope peptide and the MHC allele. An IC_50_ value of ≤500nM is considered for CD8+ T cell epitopes whereas ≤1000nM is used for CD4+ T cell epitopes (Nielsen, Lundegaard et al. 2003).

### Bovine BoLA class I- and II-restricted epitope prediction

Like MHC-specific T cell epitopes in murine system, CD8+ and CD4+ T cell epitopes in the NS1, NS2 and NS3 proteins of BTV1 serotype were predicted for class I and II alleles of BoLA. For this, host species “cow” was selected for the prediction of CD8+ T cell epitopes in IEDB server (http://tools.iedb.org/mhci/) (Vita, Mahajan et al. 2019) The most frequent class I alleles of BoLA-1*02301, BoLA-2*01201, BoLA-3*00201, BoLA-4*02401, BoLA-6*01301 and BoLA-6*01302 were considered for predicting CD8+ T cell epitopes based on previous studies (Pandya, Rasmussen et al. 2015, Svitek 2015). IC_50_ value of ≤500nM was used as a threshold for the prediction of BoLA class I allele-specific CD8+ T cell epitopes. For the prediction of BoLA II CD4+ T cell epitopes, NetBoLAIIPan 1.0 (https://services.healthtech.dtu.dk/service.php? NetBoLAIIpan-1.0) tool was used at default settings (Fisch, Reynisson et al. 2021). This tool generates the possible predicted T cell epitopes in 15 AA length with different %Rank EL scores, which is a threshold parameter for the prediction of CD4+ T cell epitopes. The peptides with %Rank EL scores of less than 1 are considered as epitopes according to the tool (Fisch, Reynisson et al. 2021). For the prediction of CD4+ T cell epitopes, all the class II alleles of BoLA-BoLA-DRB3_0101, BoLA-DRB3_1001, BoLADRB3_1101, BoLA-DRB3_1201, BoLA-DRB3_1501, BoLADRB3_1601 and BoLADRB3_2002 (these alleles well characterized using both immunoinformatics and advanced mass spectroscopy methods) deposited in the tool were selected for predicting CD4+ T cell epitopes.

### Epitope conservation

All predicted T cell epitopes, including both CD8+ and CD4+ epitopes, underwent thorough verification for their conservation across all BTV serotypes. To accomplish this, we scrutinized the predicted epitopes located within the NS1, NS2, and NS3 proteins of BTV1 by cross-referencing them with the amino acid alignment files previously generated using MEGA-X software. Our analysis focused on ascertaining the presence and degree of conservation of these BTV1 epitopes within the alignment files encompassing NS1, NS2, and NS3 proteins across all 24 serotypes. The results of this conservation assessment are presented and discussed comprehensively within this study.

### Vaccine design

Highly conserved CD8+ and CD4+ T cell epitopes were used for the design of a multi-epitope broad-spectrum BTV vaccine. Firstly, the T cell epitopes were screened for the antigenicity, allergenicity, toxicity and IFNγ-inducing abilities with the help of VaxiJen v2.0 (http://www.ddg-pharmfac.net/vaxijen/VaxiJen/VaxiJen.html) (Doytchinova and Flower 2007), AllerTOP v2.0 (https://www.ddg-pharmfac.net/AllerTOP/) (Dimitrov, Bangov et al. 2014), ToxinPred (http://crdd.osdd.net/raghava/toxinpred/) (Gupta, Kapoor et al. 2013) and IFNepitope predict (https://webs.iiitd.edu.in/raghava/ifnepitope/application.php) (Dhanda, Vir et al. 2013) web servers. Finally, the antigenic, non-allergic, non-toxic, and IFNγ-inducing conserved T cell epitopes were used in vaccine design. The CD8+ and CD4+ T cell epitopes were joined together with the help of ‘AAY’ and ‘GPGPG’ linkers. An adjuvant sequence was attached to the vaccine sequence at the N-terminal end with the helper of ‘EAAK’ linker. The adjuvants used for the vaccine design are beta-defensin as toll-like receptor (TLR) 3 agonist and 50S ribosome subunit sequence as TLR4 agonist.

### Antigenicity, allergenicity, solubility, and physicochemical properties of the vaccine constructs

We designed a total of 4 vaccine constructs which included 2 mouse (each with TLR3 agonist or TLR4 agonist adjuvant) and 2 bovine vaccines (each with TLR3 agonist or TLR4 agonist adjuvant).

These vaccines were then evaluated for their antigenicity and allergenicity as above. The solubility and physicochemical properties, including molecular weight, theoretical isoelectric point (pI), instability index (II), aliphatic index (AI), and grand average of hydropathicity index (GRAVY), of the vaccine were predicted using Protein-Sol (http://protein-sol.manchester.ac.uk/) (Hebditch, Carballo-Amador et al. 2017) and ProtParam tool (https://web.expasy.org/protparam/), respectively.

### Structure modeling and evaluation

The three-dimensional structure of the vaccine constructs designed in the study were predicted using Robetta server (https://robetta.bakerlab.org/) (Kim 2004). The top model of all the four vaccine constructs were further refined for docking studies. Subsequently, we refined the modeled 3D structures using the GalaxyWEB server (Ko, Park et al. 2012). The quality of the modeled structures was assessed by analyzing phi (ϕ) and psi (Ψ) torsion angles using PROCHECK (R. A. Laskowski 1993) through the protein structure verification server (PSVS) (https://montelionelab.chem.rpi.edu/PSVS/PSVS/).

### Preparation of the receptors and molecular docking

The ability of the receptors, TLR3 and TLR4, to interact with vaccine constructs were analyzed by molecular docking. These two TLRs are well known to elicit antiviral activity (Suh 2009). The 3D coordinates of mouse TLR3 (PDB ID: 2A0Z) and mouse TLR4 (PDB ID: 4G8A) proteins were obtained from the RCSB-PDB (Berman, Westbrook et al. 2000) in pdb format. The bovine TLR3 (Q5TJ59) and TLR4 (Q9GL65) were obtained from the alfa-fold database. Before docking, these receptor molecules were prepared with the help of MGLTools (Morris, Huey et al. 2009) and the UCSF Chimera package (Pettersen, Goddard et al. 2004). For molecular docking studies, Cluspro server (Yan, Tao et al. 2020) was utilized. In the docking process, the TLR3 and TLR4 proteins served as receptors (chain A), while the designed vaccine constructs were used as ligand molecules (chain B). The top-ranked docking complexes were selected and the binding affinity between the receptor and ligand in the docked complexes was determined with PRODIGY server (Li C. Xue 2016). The PDBsum server (Laskowski, Jablonska et al. 2018) was utilized to generate the molecular interaction graphs.

### Molecular dynamics simulations

A molecular dynamics simulations (MDS) framework was established for all top four docking complexes to investigate their stability at the atomic level. The same protocol was applied to every complex, and state-of-the-art MDS was conducted for 100 nanoseconds using the AMBER99SB-ILDN protein, nucleic AMBER94 force field (Lindorff-Larsen, Piana et al. 2010) embedded in the GROMACS 2023 package (Abraham 2015) installed on a high-performance computing (HPC) system. The docking complexes were solvated using the transferable intermolecular potential 3P (TIP3P) water model. To achieve neutrality, we employed LINear Constraint Solver (LINCS) constraint algorithms for energy minimization in all systems (Hess 1997). Moving on, the docking complexes underwent equilibration using the NVT ensemble at 300K and the NPT ensemble with the Parinello-Rahman barostat (Parrinello 1981) coupling ensembles. Subsequently, a 100ns MDS was conducted for all four docking complexes. After the MDS, we analyzed the trajectories and generated plots using various modules available in the GROMACS package.

## RESULTS

### Differing conservation levels across non-structural proteins

We conducted a thorough analysis of amino acid sequence conservation within the NS1, NS2, and NS3 proteins, employing multiple sequence alignment techniques to assess the extent of conservation. This analysis aimed to shed light on the potential presence of conserved epitopes. After the alignment of amino acid sequences, molecular data files in MEGA format with an extension ‘.meg’ or ‘.mega’ were opened, and the amino acid polymorphism was identified in each alignment file of NS1, NS2 and NS3 proteins. Our findings revealed that among the 24 BTV serotypes, the NS1, NS2, and NS3 proteins exhibited varying degrees of amino acid sequence conservation. Notably, NS1 emerged as the most highly conserved with an impressive amino acid sequence identity homology of 83.51%, followed by NS3 with 83.40% and NS2 with 73.66% sequence identities (**Figure 1A**). These results strongly suggest that these proteins might harbor a higher abundance of conserved T cell epitopes.

**Figure 1.**
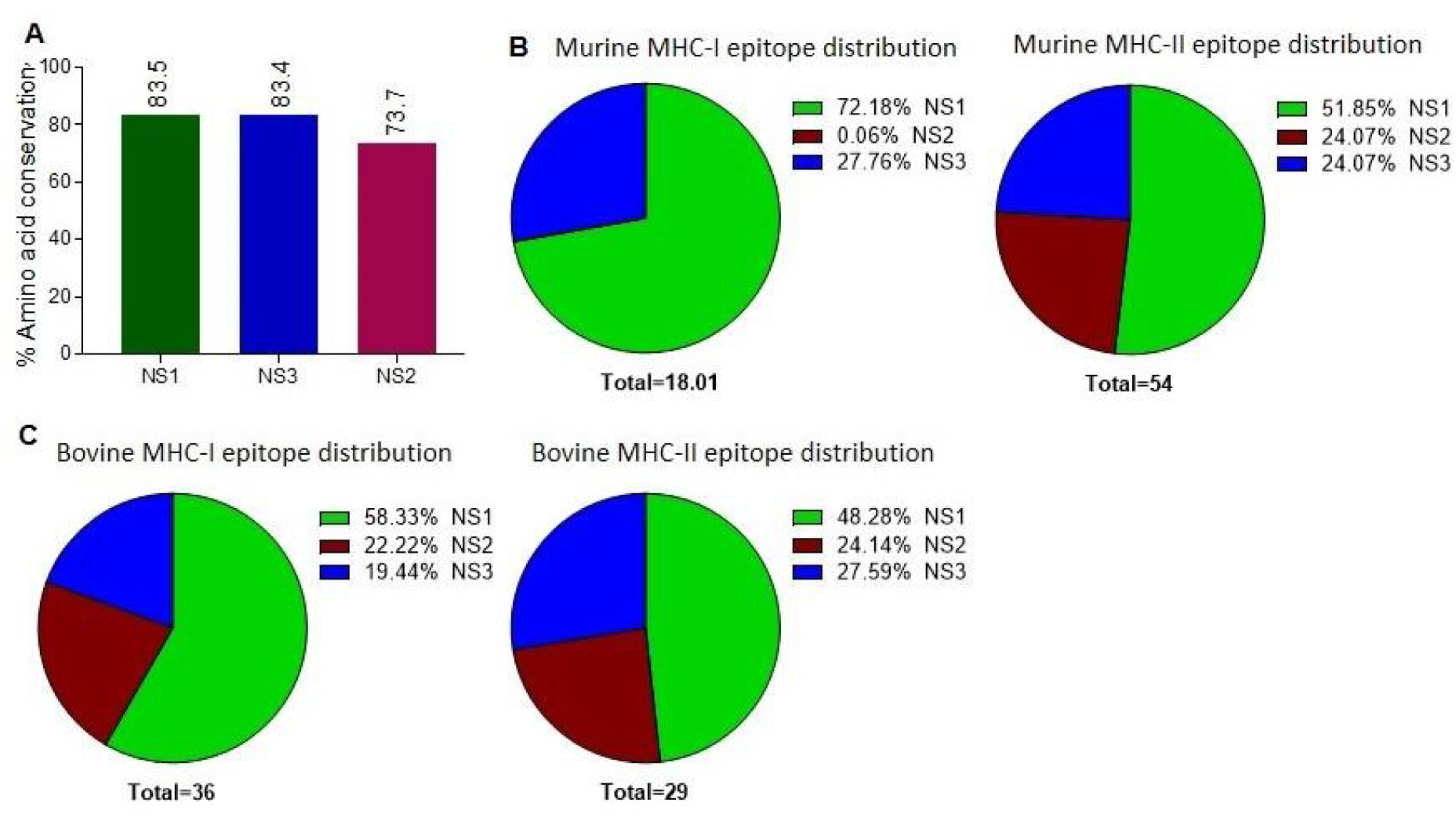
Amino Acid Sequence Conservation and T Cell Epitope Distribution in Non-Structural BTV Proteins. **A.** Amino acid sequence conservation within the non-structural proteins, highlighting pronounced conservation in NS1, NS2, and NS3 proteins. **B-C.** Distribution of murine (B) and bovine (C) CD8+ and CD4+ T cell epitopes across the NS proteins.

### Frequencies of T cell epitopes recognized by murine and bovine systems

Mouse models are increasingly becoming valuable for testing T-cell-mediated immunity of vaccines against BT (Calvo-Pinilla 2009, Martinelle, Dal Pozzo et al. 2018, Saminathan, Singh et al. 2020). Particularly, C57BL/6 background mouse model has been well established for studying the pathogenesis and vaccine development for BTV (Umeshappa, Singh et al. 2010, Umeshappa, Singh et al. 2011, Umeshappa, Singh et al. 2011, Potter 2019). Hence, we used their MHC class I H2-D^b^ and H2-K^b^, and class II H2-IA^b^ haplotypes for predicting CD8+ and CD4+ T cell epitopes in NS1, NS2 and NS3 proteins of BTV. Processing and presentation of antigenic epitopes to T cells are key events in the development of antiviral immune response. The immunogenicity of an antigen is significantly influenced by the affinity of interaction between T-cell receptors (TCRs), epitopes, and MHC complex, particularly for viral-specific CD8+ and CD4+ T cells (Sette 1994, Paul, Weiskopf et al. 2013). Previous reports indicate that an IC_50_ value of less than 500nM serves as a threshold affinity between the epitope and class I MHC molecules (Sette 1994, Paul, Weiskopf et al. 2013). Conversely, the majority of experimentally verified CD4+ T cell epitopes exhibit an affinity of less than 1000nM with class II MHC/HLA molecules (Southwood 1998, Sidney, Steen et al. 2010).

Based on these criteria, a total of 18 CD8+ T cell epitopes were predicted for mouse system, which includes 13 (72.18%) and 5 (27.76%) CD8+ T cell epitopes in the NS1 and NS3 proteins [**Figure 1B, Supplementary Table 1 (S1**)]. Surprisingly, CD8+ T cell epitopes were not observed in NS2 protein. Similarly, a total of 28 (51.85%), 13 (24.07%), 13 (24.07%) CD4+ T cell epitopes (a total of 54) were identified in the NS1, NS2 and NS3 proteins, respectively (**Figure 1C, Table S2**).

Like BTV’s natural host sheep (ovine), bovines, such as cattle, are also significant hosts and are closely related to Ovine in genome, anatomy, immunology, and physiology. Since tools are unavailable for predicting T cell epitopes for Ovine MHC alleles, we used only BoLA I and II alleles, which would also give an idea about conserved CD8+ and CD4+ T cell epitope presentation by ovine immune system. The IEDB contain information for class I BoLA alleles and has hundreds of BoLA class I alleles, which are very difficult to analyze for CD8+ T cell epitope prediction. Hence, we considered MHC class I BoLA alleles, BoLA-BoLA-6*01301, BoLA-6*01302, BoLA-2*1201, BoLA-4*02401, BoLA-3*00101 and BoLA-1*02301, which are more prevalent in bovine population and have been previously used for the prediction of BoLA-restricted class I epitopes for foot-and-mouth virus, whose infection resembles BT in the initial stages of infection (Williamson S 2008). For BoLA I-specific CD8+ T cell epitope prediction, here also we chose IC_50_ value of ≤ 500nM and used the “MHCpan BA 4.1” module in the IEDB server. We obtained a total of 36 CD8+ T cell epitopes, which includes 21 (58.33%), 8 (22.22%), 7 (19.44%) CD8+ T cell epitopes (pooled values of all 6 BoLA class I alleles) in the NS1, NS2 and NS3 proteins of BTV-1 serotype (**Figure 1D, Table S3**).

Similarly, for predicting CD4+ T cell epitopes, we used highly prevalent class II BoLA alleles in the bovine population, BoLA-BoLA-DRB3_0101, BoLA-DRB3_1001, BoLA-DRB3_1101, BoLA-DRB3_1201, BoLADRB3_1501, BoLADRB3_1601 and BoLA-DRB3_2002 (Fisch, Reynisson et al. 2021). We considered the recommended cut-off score ≤ 1.0 %Rank EL and used the tool NetBoLAIIPan 1.0 (Fisch, Reynisson et al. 2021). Here, we obtained a total of 66 CD4+ T cell epitopes (pooled values of all 7 BoLA class II alleles) with 14 (48.28%), 7 (24.14%), 8 (27.59%) presents in the NS1, NS2 and NS3 proteins of BTV-1 serotype, respectively (**Figure 1E, Table S4**).

### Conserved T cell epitope presentation by a laboratory murine model

Through sequence alignment, we found highly conserved T cell epitopes in NS1, NS2, and NS3 of all the 24 BTV serotypes (**Figure 2**). One CD8+ T cell epitope H2-K^b^-KHFNRYASM was found to be 100 % conserved in NS1 and two epitopes (AAFASYAEA and KAMSNTTGA) in NS3 (**Figure 2A, B; Table S1**). Though all the epitopes in these proteins are not conserved 100%, a majority of the amino acids in those epitopes are conserved with at least 60% conservation, which is cut-off value to be considered for an epitope to be cross-reactive (Tengvall 2019, Girdhar, Huang et al. 2022). In NS1, the amino acid conservation in the epitopes is 88.88% (9), 77.77% (1) and 66.66% (1) (**Table S1**). Similarly, NS3 protein have 2 CD8+ T cell epitopes each with 88.88% and 66.66% of the conserved amino acids.

**Figure 2.**
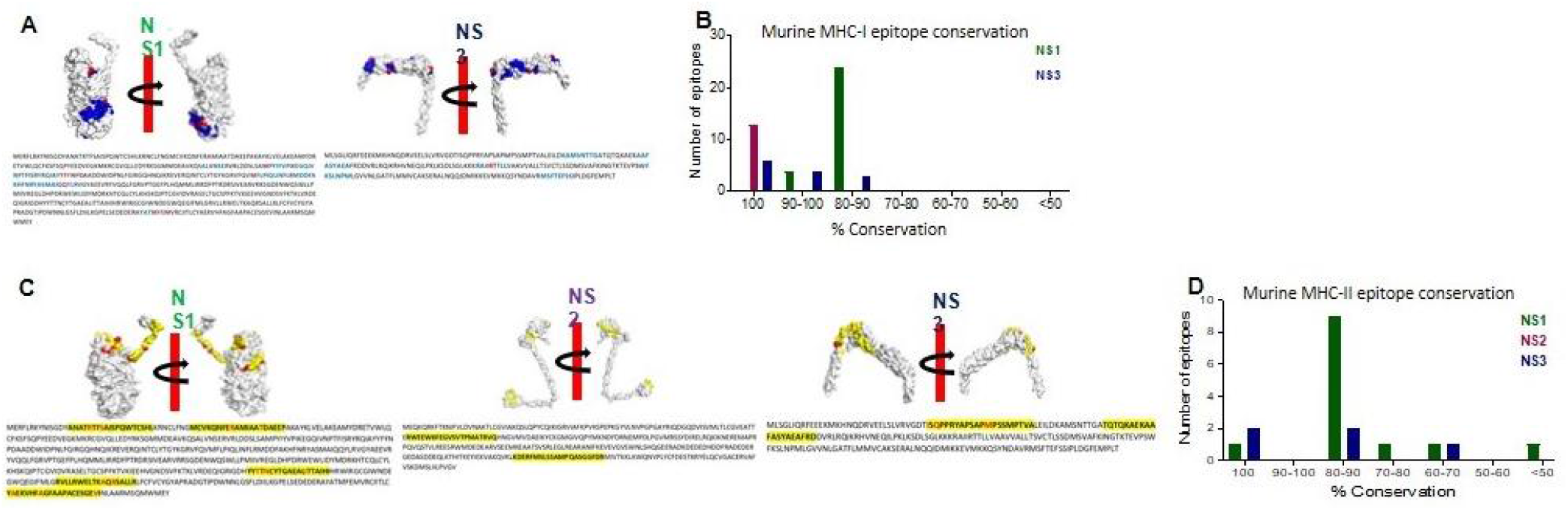
Conserved CD8+ and CD4+ T Cell Epitope Presentation in a Laboratory Murine Model. **A.** Protein structures depicting MHC class I epitope conservation (highlighted in blue) and variations in amino acid residues within the epitopes (highlighted in red). *In silico* protein structure predictions were generated using the Robetta server, with both front and back views displayed with a 180-degree rotation. **B.** Bar chart illustrating the extent of amino acid conservation among MHC class I-restricted CD8+ T cell epitopes within NS1 and NS3. **C.** Protein structures showcasing MHC class II epitope conservation (highlighted in blue) and variations in amino acid residues within the epitopes (highlighted in red). The protein structures were generated *in silico* through the Robetta server, and both front and back views are presented with a 180-degree rotation. **D.** Bar chart presenting the degree of amino acid conservation among MHC class II-restricted CD4+ T cell epitopes within NS1, NS2, and NS3.

Within the conserved CD4+ T cell epitopes, it is noteworthy that the NS1 protein, characterized by its remarkable ∼83.5% sequence identity, contains a substantial proportion of these epitopes. Specifically, we identified 4 epitopes with an impressive 93.33% sequence identity, along with 18 epitopes exhibiting an 86.66% sequence identity, and an additional 6 epitopes with approximately 80% sequence identity (**Figure 2C, D; Table S2**). Interestingly, all the CD4+ T cell epitopes in NS2 protein are conserved with 100% sequence identity (**Table S2**). In NS3 protein, 6 CD4+ T cell epitopes out of 13 were found as 100% conserved ones and the remaining 4 CD4+ T cell epitopes have 93.33% (#1), 86.66% (#2) and 80% (#1) sequence identity (**Table S2**).

### Distribution of conserved CD8+ and CD4+ T cell epitopes in Bovines

Similarly, in the case of BoLA MHC-restricted conserved T cell epitopes, we detected a discernible trend of CD8+ T cell epitope conservation within the NS1, NS2, and NS3 proteins (**Figure 3A, B**). The NS1 protein contain highest number of 100% conserved CD8+ T cell epitopes, AMYDRETVW, KHFNRYASM and RKYNISGDY. Only one CD8+ T cell epitope is least conserved with 55.55%. Among the remaining 17 CD8+ T cell epitopes, 9 are with 88.88%, 7 with 77.77% and 1 is with 66.66% sequence identify (**Figure 3A; Table S3**). Only 1 CD8+ T cell epitope in NS2 is 100% conserved among all the 24 BTV serotypes. 6 of the predicted CD8+ T cell epitopes contain 88.88% of the amino acid conservation in the NS2 protein. Whereas only 1 CD8+ T cell epitope is least conserved in NS2 protein with 55.55%. In NS3 protein, 1 CD8+ T cell epitope is highly conserved with 100% conservation and the remaining CD8+ T cell epitopes contain > 65% of the amino acid conservation i.e., with 88.88%, 2 with 77.77% and 2 with 66.66% sequence identity. Interestingly, the NS3 protein also contain CD4+ T cell epitope, LRQIKRHVNEQILPK, which is 100% conserved.

**Figure 3.**
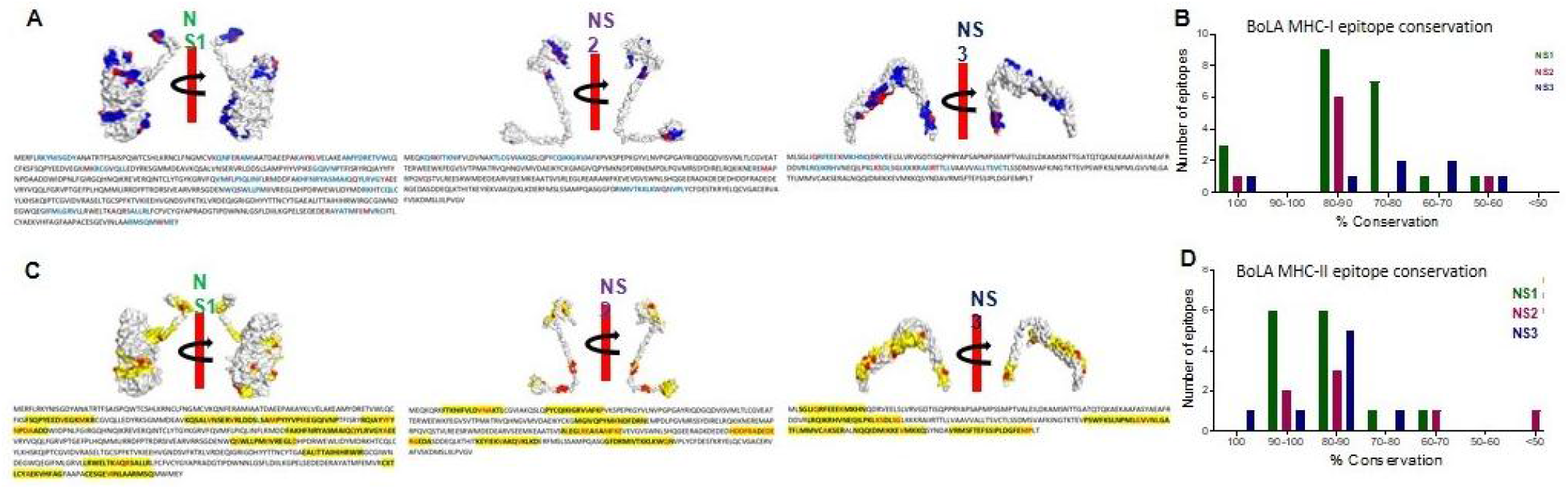
Conserved CD8+ and CD4+ T Cell Epitope Presentation by Bovine Hosts. **A.** Protein structures illustrating MHC class I epitope conservation (highlighted in blue) and the variations in amino acid residues within the epitopes (highlighted in red). *In silico* protein structure predictions were generated using the Robetta server, with both front and back views displayed along with a 180-degree rotation. **B.** Bar chart presenting the extent of amino acid conservation among MHC class I-restricted CD8+ T cell epitopes within NS1, NS2, and NS3. **C.** Protein structures demonstrating MHC class II epitope conservation (highlighted in blue) and variations in amino acid residues within the epitopes (highlighted in red). The protein structures were produced *in silico* through the Robetta server, and both front and back views are presented with a 180-degree rotation. **D.** Bar chart depicting the degree of amino acid conservation among MHC class II-restricted CD4+ T cell epitopes within NS1, NS2, and NS3.

Much like the conservation observed in CD8+ T cell epitopes, a similar pattern emerged in CD4+ T cell epitopes within the NS1, NS2, and NS3 proteins (**Figure 3C, D**). Among these, NS3 exhibited the highest conservation of CD4+ T cell epitopes, reaching 100%, followed by NS1 and NS2 (**Figure 3D; Table S4**). Within NS1, 12 out of 14 predicted CD4+ T cell epitopes were conserved, with sequence identities of 93.33% and 86.66% and the remaining 2 epitopes have sequence identities of 73.33% and 66.66% (**Figure 3C; Table S4**). Similarly, in NS2, all CD4+ T cell epitopes demonstrated conservation, with over 60% sequence identity. This included 93.33% for 2 epitopes, 86.66% for 1, 80% for 2, and 60% for 1, with just one epitope showing a lower conservation at 26.66% (**Figure 3C; Table S4**). Notably, NS3, being highly conserved, harbored CD4+ T cell epitopes with more than 73% sequence conservation, with an identity of 100% for 1 epitope, 93.33% for 1, 86.66% for 2, 80% for 3, and 73.33% for 1 (**Figure 3C; Table S4**).

### Non-structural proteins are hotspots for conserved epitopes in both murine & bovine systems

NS proteins, intriguingly, serve as prominent source for conserved epitopes in both murine and bovine systems (**Figure 4A**). A substantial number of CD8+ and CD4+ T cell epitopes, with a minimum amino acid conservation of over 60%, are predominantly situated within NS1, NS2, and NS3 proteins, respectively. It is noteworthy that in the mouse system, there were no corresponding epitopes found in the NS2 protein for MHC class I alleles. What is remarkable is that this pattern holds true in both the mouse and bovine systems. These findings underscore the significance of NS proteins as valuable candidates for inclusion in considerations for the development of BT vaccines.

**Figure 4.**
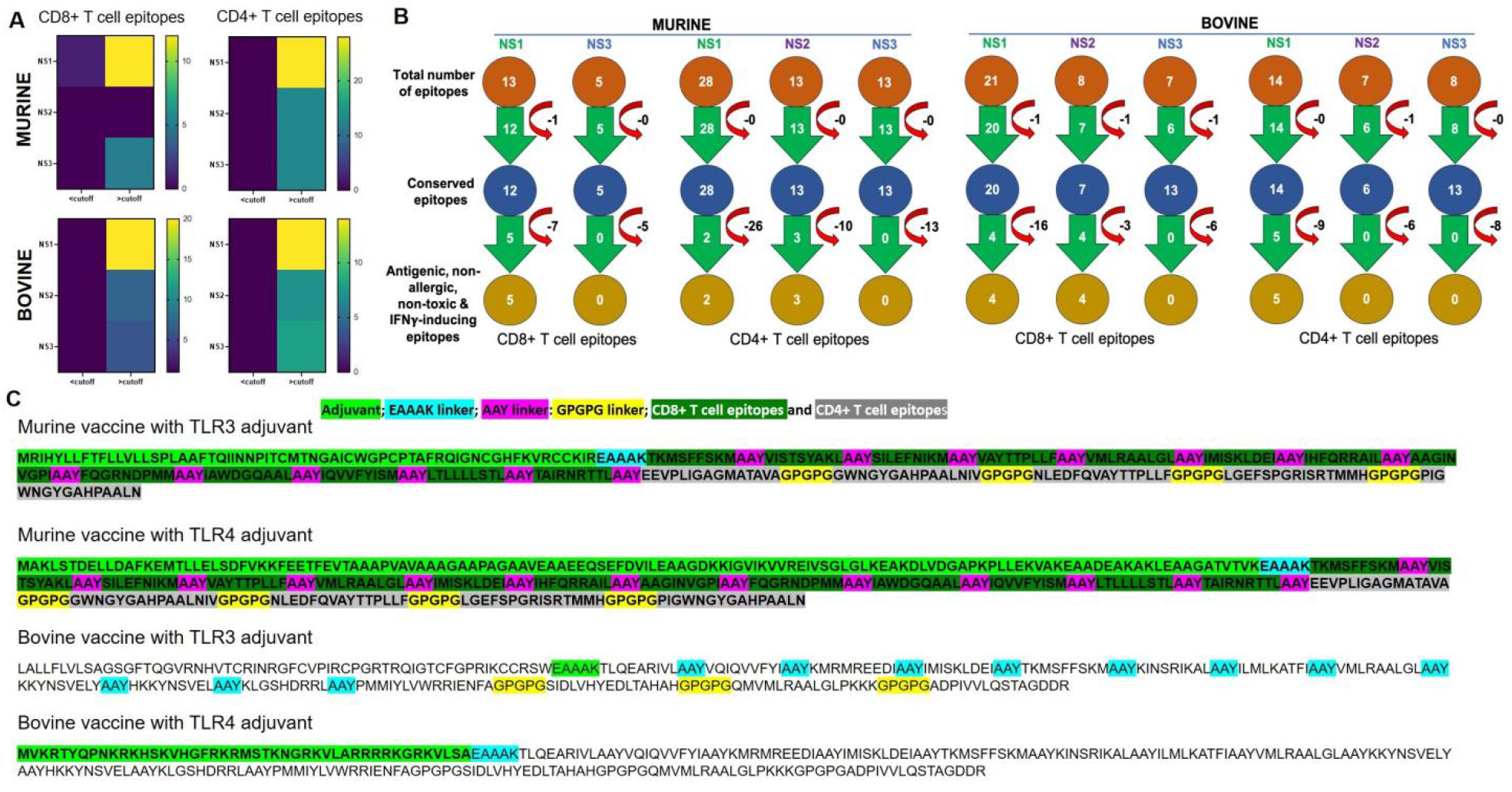
Conserved Epitopes and Multi-Epitope Vaccine Design for Bluetongue Virus. **A.** Non-Structural Proteins Serve as Hotspots for Conserved Epitopes in Both Murine and Bovine Systems. Heat maps demonstrate the increased prevalence of CD4+ and CD8+ T cell epitopes, particularly within the NS1 protein. **B.** Schematic representation of the screening process for identifying highly conserved immunodominant epitopes, used in the development of a multi-epitope pan-BTV vaccine. **C.** Design of the multi-epitope pan-BTV vaccine for both murine and bovine systems. The adjuvant is attached at the N-terminal of the vaccine sequence using an EAAK linker, while CD8+ and CD4+ T cell epitopes are connected with AAY and GPGPG linkers, respectively.

### Screening of conserved epitopes and *in silico* broad-spectrum vaccine formulation

Next, we screened all the conserved epitopes for their antigenicity, allergenicity, toxicity, and IFNγ-inducing abilities. After screening for these criteria, we obtained a total of 5 MHC I-specific CD8+ T cell epitopes in NS1 and none in NS2 and NS3 proteins and a total of 2 MHC II-specific CD4+ T cell epitopes in NS1, 3 in NS2 and none in NS3 for murine vaccine development **(Figure 4B)**. For bovine vaccine development, we obtained a total of 4 BoLA I-specific CD8+ T cell epitopes in NS1, 4 in NS2 and none in NS3 and a total of 5 BoLA I-specific CD8+ T cell epitopes in NS1 and none in NS2 and NS3 proteins. These filtered epitopes are capable of mounting robust anti-viral T cell response (antigenic and IFN γ -inducing) and do not cause any toxicity, autoreactivity and allergenicity *in vivo*. Hence, they were used for designing *in silico* multi-epitope broad-spectrum BTV vaccine formulation as illustrated in method section (**Figure 4C**). Briefly, the conserved T cell epitopes were joined together with the help of linkers, and then TLR3 and TLR4 agonist adjuvants were added to the N-terminus region of the CTL epitopes to boost the immunogenicity (Pyasi, Sharma et al. 2021). A total of 2 constructs were designed for mouse and 2 for bovine.

### Antigenicity, allergenicity, solubility, and physicochemical properties of the vaccine constructs

Later, we evaluated the vaccine constructs *in silico* to ensure their safety and immunogenicity. Based on the antigenicity analysis (**Table 1**), all the four vaccine constructs are considered as probable antigens. Similarly, the allergenicity and solubility analyses indicated that all four vaccine constructs are non-allergen and safe for vaccine development and good soluble (**Table 1**).

**Table 1.** Immunological and physicochemical properties of the designed vaccine constructs.

| Vaccine | Antigenicity | Allergenicity | Soluble | Molecular weight | Isoelectric point | Instability index | Aliphatic index | GRAVY |
| --- | --- | --- | --- | --- | --- | --- | --- | --- |
| Mouse-TLR3 | Antigenic (0.6839) | Non-allergen | Soluble (0.395) | 24512.71 | 9.76 | 21.28 | 77.09 | 0.213 |
| Mouse-TLR4 | Antigenic (0.5900) | Non-allergen | Soluble (0.538) | 30824.54 | 5.53 | 20.52 | 83.59 | 0.11 |
| Bovine-TLR3 | Antigenic (0.6253) | Non-allergen | Soluble (0.501) | 29783.16 | 9.78 | 34.3 | 95.54 | 0.134 |
| Bovine-TLR4 | Antigenic (0.6098) | Non-allergen | Soluble (0.727) | 27111.99 | 10.21 | 37.42 | 90.33 | -0.208 |

The physicochemical properties evaluation showed that mouse-TLR3, mouse-TLR4, bovine-TLR3 and bovine-TLR4 vaccine constructs have the molecular weights, 24512.71, 30824.54, 29783.16 and 27111.99, and theoretical isoelectric point (pI) values, 9.76, 5.53, 9.78 and 10.21, respectively (**Table 1**). Further, these constructs have the instability indices of 21.28, 20.52, 34.3 and 37.42, respectively, indicating that all the four vaccine constructs are stable. In addition, these constructs have higher aliphatic index values, 77.09, 83.59, 95.54 and 90.33, respectively, suggesting their thermostability. Finally, GRAVY scores indicates that mouse-TLR3, mouse-TLR4, and bovine-TLR3 constructs are polar in nature (0.213, 0.11, and 0.134, respectively) and bovine-TLR4 vaccine construct is non-polar in nature (-0.208) (**Table 1**).

### Structural modeling and evaluation of the vaccine constructs

In this study, we utilized the revolutionary method for the structure modeling of the designed vaccine constructs **(Figure 5A)**. Molecular modeling, is one of the principal and accurate methods for protein structure modeling based on its amino acid sequence (Muhammed and Aki-Yalcin 2019). Structural evaluations of the 3D structure models of vaccine constructs using Ramachandran plot calculations are a popular approach and have frequently been utilized in several recent studies (Aiman, Alhamhoom et al. 2022, Samad, Meghla et al. 2022). As per the general criteria of Ramachandran plot analysis, a model with ∼90% of residues in the most favored regions are considered of good quality. The modeled 3D structure models were evaluated by calculating their phi (ϕ) and psi (Ψ) torsion angles using Ramachandran plot analysis. As evident from **Figure 5B**, the generated Ramachandran plots statistics for the mouse vaccine-TLR3, mouse vaccine-TLR4, bovine vaccine-TLR3 and bovine vaccine-TLR4 modeled structures revealed a total of 86.9, 92.7, 90.8 and 96% of residues, respectively, were found in favorable regions, and 10.4, 6.1,7.2 and 3% were in additionally allowed regions, respectively **(Supplementary Table 6)** and 0.5, 0, 1.4 and 0% were found in the generously allowed regions and no single residue was found scattered in disallowed regions of the Ramachandran plots, confirming the excellent quality of the modeled 3D structures of the vaccine constructs. The structural evaluation of the structural models revealed that the predicted structures of the vaccine constructs are of good quality and suitable for molecular docking.

**Figure 5.**
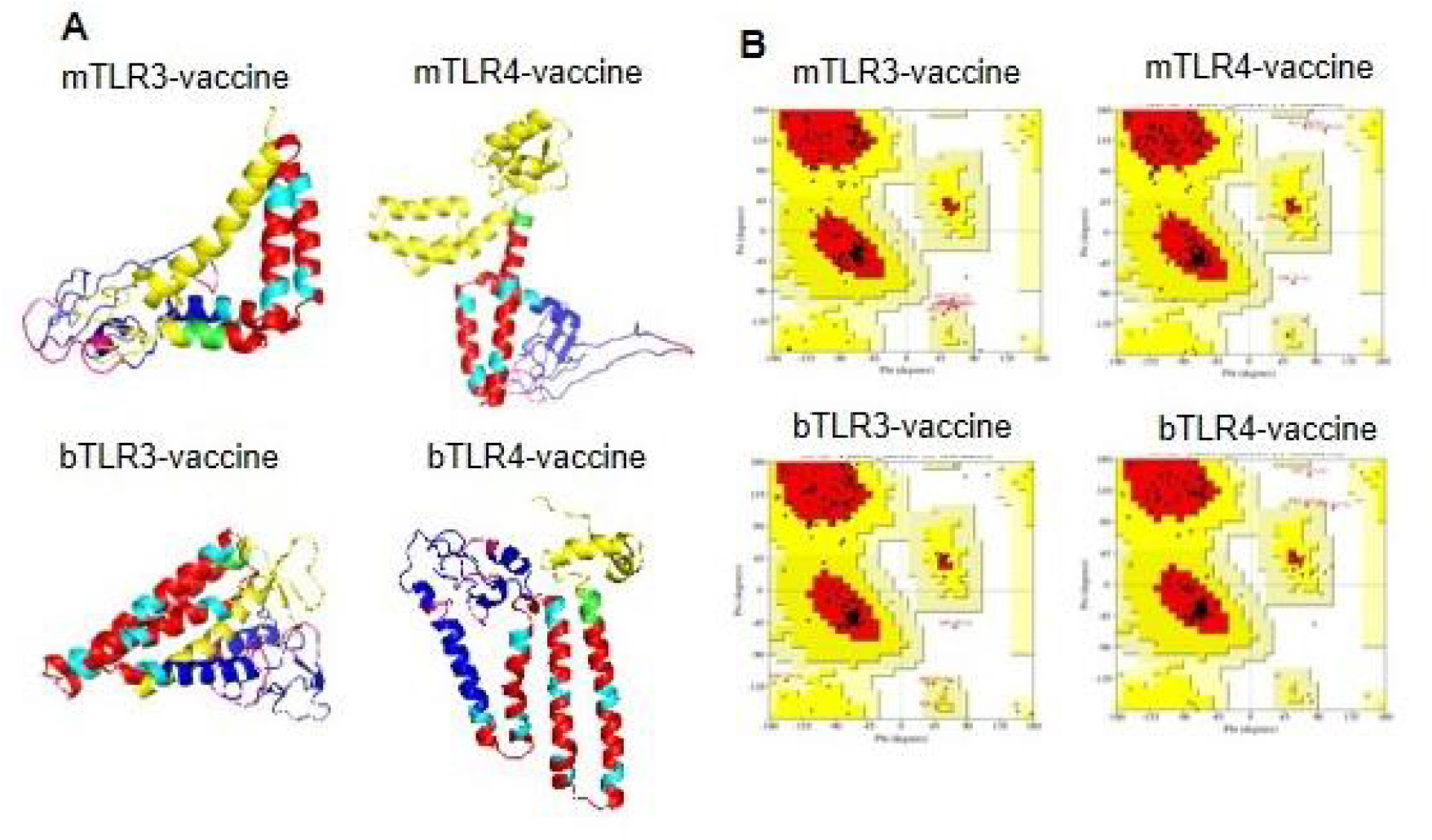
Modeling and Evaluation of Three-Dimensional Structures of the Vaccine Constructs. **A.** Three-dimensional structures of the vaccine constructs obtained through protein modeling. **B.** Ramachandran plots illustrating the protein models of vaccine constructs obtained through protein modeling.

### Molecular docking

Protein-protein docking has been established as one of the popular approaches for the prediction of molecular interaction patterns between the toll like receptor receptor (TLR) molecules and vaccine construct (Vakser 2014). To determine the molecular interaction between designed vaccine constructs and TLRs, we have performed a molecular docking. The mouse and bovine vaccine constructs with TLR3 antagonist were docked against TRL3, while two other constructs with TLR4 antagonist were docked against TLR4 using the HDOCK platform (**Figure 6A, B**). During the docking analysis, it was observed that the vaccine constructs for mouse-TLR3, mouse-TLR4, bovine-TLR3, and bovine-TLR4 exhibited the Gibbs free energy (**ΔG) values** of -–19.3 Kcal/mol, – 18.4 Kcal/mol, –12.6 Kcal/mol and –16.3 Kcal/mol respectively **(Figure 6C)**. Out of these four docked vaccine constructs, the mouse-TLR3 ranked as the top interacting construct against its receptor protein, based on the calculated higher negative docking score. The docking complexes of these vaccine constructs with TLRs exhibited several molecular interactions including hydrogen bonds, salt bridges, and non-bonded contacts.

**Figure 6.**
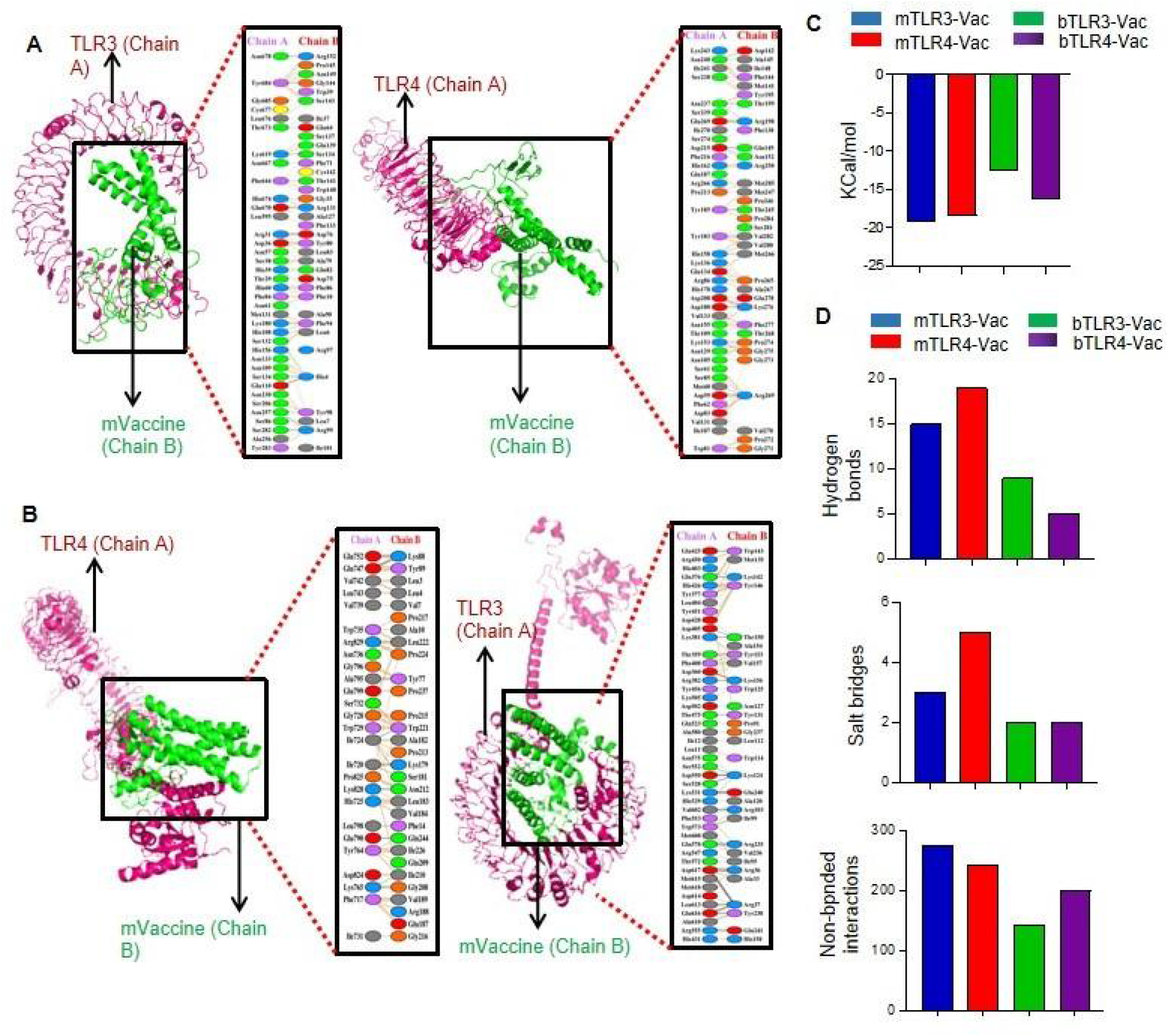
Interaction of the Vaccine Constructs with Mouse and Bovine TLRs. **A.** Molecular interactions of the mouse vaccine constructs (chain B) with their respective TLR3 (left) and TLR4 (right), visualized in both 3-D and 2-D views. **B.** Molecular interactions of the bovine vaccine constructs (chain B) with their corresponding TLR3 (left) and TLR4 (right), represented in both 3-D and 2-D views. **C.** Docking scores for the murine and bovine vaccine docked complexes, including mTLR3-Vac, mTLR4-Vac, mouse vaccine-TLR3, bTLR3-Vac, and bTLR4-Vac. D. Bar chart illustrating the total number of salt bridges, hydrogen bonds, and non-bonded interactions between the vaccine constructs and TLRs.

The 2-D molecular interactions were visualized using PDBSum, revealing that the docking complex of the mouse vaccine and TLR3 was stabilized by 15 hydrogen bonds, 3 salt bridges, and 276 non-bonded interactions **(Figure 6A and 6D)**. The mouse vaccine-TLR4 complex formed 19 hydrogen bonds, 05 salt bridges, and 244 non-bonded contacts (**Figure 6A and 6D**). The bovine vaccine-TLR3 complex formed the 09 hydrogen bonds, 02 salt bridges, and 144 non-bonded contacts (**Figure 6B and 6D**). In the complex of bovine vaccine and TLR4, 05 hydrogen bonds, 02 salt bridges, and 200 non-bonded interactions have been calculated (**Figure 6B and 6D**). Based on the molecular interaction patterns, the complex of mouse vaccine and TLR3 was found to have the second highest number of hydrogen bonds after mouse vaccine-TLR4 complex and 276 non-bonded interactions. The outcome of molecular docking is consistent with the docking results of the previous studies (Bhattacharya and Roy 2008, Sun, Sun et al. 2014, Russell, Parbhoo et al. 2018).

### Molecular dynamics simulations on 100 ns

MDS was performed to assess the stability of the docking complexes of the vaccine and receptor complexes at atomic level on 100 ns. The dynamic behaviour of the simulated systems was investigated using different functions including root mean square deviation (RMSD) and root mean square fluctuation (RMSF) available in GROMACS.

RMSD plot analysis is a well-established method to measure the changes in the protein structure during MDS. In the present study, calculated RMSDs of the vaccine and receptors docking complexes were graphically investigated to assess the stability at the atomic level. As evident from **Figure 7A**, vaccine and receptor docking complexes demonstrated a constant range of stability throughout the simulation on 100ns with a range between ∼0.52 and ∼2.15nm. The average RMSD values for the mouse vaccine-TLR3, mouse vaccine-TLR4, bovine vaccine-TLR3 and bovine vaccine-TLR4 were 0.52, 0.65, 2.15 and 1.9 respectively. As illustrated in **Figure 7A**, docking complexes of mouse vaccine constructs showed a small level of fluctuations at the starting point between 5 to 20ns, while bovine vaccine constructs demonstrated the multiple fluctuations throughout the MD simulations on 100ns.

**Figure 7.**
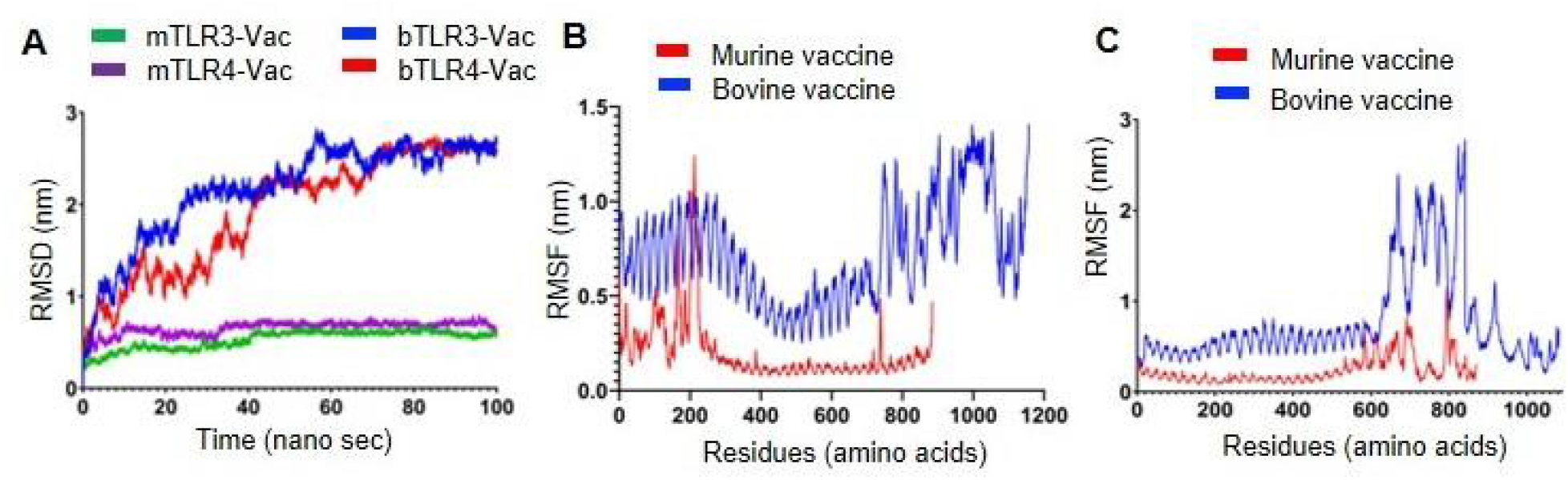
RMSD and RMSF Plots of the Docking Complexes: **A.** Line diagram displaying the calculated RMSD complexes between the mouse and bovine vaccines and TLRs. **B.** Line diagram illustrating the RMSF of the docking complexes between mouse and bovine vaccine constructs and TLR3. **C.** Line diagram illustrating the RMSF of the docking complexes between mouse and bovine vaccine constructs and TLR4.

The complex of bovine vaccine-TLR3 (blue) showed three major fluctuations, the first fluctuation was observed between ∼05 to 20ns, the second between ∼40 to 60ns, while the third major fluctuation was noted between ∼65 to 85ns. After, 85ns, the complex of bovine vaccine-TLR3 (blue) reflected the stability up to around 100ns time scale on ∼2.90nm. The complex of bovine vaccine-TLR4 (red) presented as the other most fluctuated; this docking complex also showed three major fluctuations between ∼05 to20, ∼25to 40ns, and ∼55-75ns. After 80 ns, the complex of bovine vaccine-TLR4 (red) showed stability up to 100ns on ∼2.90nm. Two complexes represent mouse vaccine constructs, mouse vaccine-TLR3 (green) and mouse vaccine-TLR4 (purple) exhibited an almost similar type of stability pattern throughout the simulation on a 100ns time scale. No major fluctuations have been observed in these complexes. Based on the RMSD plot analysis, it can be concluded that calculated backbone RMSDs of docking complexes indicated minimally conformational changes and vaccine and receptor complexes are stable at the atomic level.

The RMSF plot analysis was performed for the measurement of individual residue flexibility on a 100ns time scale. The higher RMSF value depicts better flexibility, while the lower RMSF value indicates correct structure regions in the docking complexes of the vaccine and receptors (56). In the present study, the RMSFs of the alpha carbon atoms of all four docking complexes were studied. All four simulated complexes namely, mouse vaccine-TLR3, and mouse vaccine-TLR4, bovine vaccine-TLR3 and bovine vaccine-TLR4 exhibited a pattern of stability with several fluctuations throughout the simulation on 100ns. The average RMSF scores of the mouse vaccine-TLR3, and mouse vaccine-TLR4 docking complexes were 0.19, 0.23, 0.72 and 0.75 respectively. It is observed from the measured RMSF plot in **Figure 7B** that the bovine vaccine-TLR3 (blue) showed the highest peak between 800 to 1150 residues on ∼1.4nm when compared to mouse vaccine-TLR3 (red). The complex of mouse vaccine-TLR3 also exhibited a higher peak around 200 residues at ∼1.3nm during simulation. **Figure 7C** indicates that complex of bovine vaccine-TLR4 (blue) and mouse vaccine-TLR4 (red) showed a similar region for a peak between 600 to 800 residues throughout the simulation on a 100 ns time scale. However, the complex of bovine vaccine-TLR4 (blue) demonstrated the highest peak on ∼2.8nm. Taken together, the few noted peaks observed in the vaccine-receptor complexes support the molecular docking results and suggest that all the vaccine constructs significantly interact with TLRs.

## DISCUSSION

Vaccination stands as the most effective strategy for combating BTV infections in ruminants, reducing susceptibility and facilitating safe animal movement from enzootic regions. Presently, globally employed vaccines against BTV include live-attenuated vaccines (LAVs) and inactivated vaccines (IVs) derived from the whole BTV (Bhanuprakash 2009, Umeshappa, Singh et al. 2010, Umeshappa, Singh et al. 2011, Ranjan 2019). These vaccines aim to stimulate host immune responses, but they are hampered by several limitations in preventing BTV spread effectively. LAVs, while highly immunogenic and offering good BTV protection, are primarily serotype-specific, with limited cross-protection capabilities (Huismans and Erasmus 1981, Sette 1994, Bhanuprakash 2009, Umeshappa, Singh et al. 2010, Umeshappa, Singh et al. 2010, Umeshappa, Singh et al. 2011, Paul, Weiskopf et al. 2013, Ranjan 2019). They also continue to circulate, contributing to genetic diversity and potential reassortment with wild-type strains. LAVs have also raised concerns related to virulence reversion, fetal malformations, and other adverse effects (Umeshappa, Singh et al. 2011). IVs, while safer than LAVs, typically confer serotype-specific and often weaker immunity (Umeshappa, Singh et al. 2011). Consequently, conventional vaccines are often serotype-specific and fall short in providing comprehensive protection against different BTV serotypes.

Some level of heterologous protection against BTV-23 was previously achieved through BTV-1 IVs, although their impact on other serotypes has not been explored extensively (Sidney, Steen et al. 2010, Umeshappa, Singh et al. 2011). Subsequently, similar instances of cross-protection were observed in various BTV vaccination scenarios. For instance, research indicated that vaccination against BTV-8 also conferred cross-protection against BTV-1 infection (Hund, Gollnick et al. 2012). Similarly, vaccination against BTV-8 was found to cross-protect against BTV-4 and BTV-1 infections (Martinelle, Dal Pozzo et al. 2018). Moreover, vaccination targeting BTV-9, -2, and -4 was shown to cross-protect against BTV-16 infection (Breard, Belbis et al. 2015). These findings together strongly suggest that BTV proteins encompass conserved epitopes capable of inducing cross-protection against serologically distinct BTV serotypes, motivating our exploration of the BTV NS proteins to identify potential conserved epitopes for the development of broad-spectrum vaccines.

Taking cues from analogous viruses with rapid evolution rates, such as influenza, we can strategize the development of a pan-BTV vaccine (Potter 2019, Evseev and Magor 2021). The presence of multiple strains of Influenza and frequent antigenic changes necessitate innovative vaccine approaches (Wang, Tang et al. 2022, Zheng, Guo et al. 2022). Studies have demonstrated that epitope-specific T cell responses and T cell clone adoptive transfers can confer cross-protection against antigenically distinct strains (Southwood 1998, Williamson S 2008, Bhanuprakash 2009, Wang 2010, Sistere-Oro, Lopez-Serrano et al. 2019). Recent research even indicates that immunogenic consensus regions could serve as the basis for universal influenza vaccines (Erasmus 1990, Jazayeri and Poh 2019, Ranjan 2019, Subbiah, Oh et al. 2022). Such strategies can be applied to pan-BTV vaccine development.

Bioinformatics tools for sequence analysis and epitope mapping can assist in identifying immunogenic consensus sequence (ICS) epitopes among BTV serotypes. In this study, we have provided computational evidence of the existence of ICS epitopes within BTV proteins, guiding vaccine development efforts. Cellular immunity plays a crucial role in providing cross-protection against different BTV serotypes (Umeshappa, Singh et al. 2010, singh 2011, Umeshappa, Singh et al. 2011, Rojas, Rodriguez-Calvo et al. 2017, Potter 2019, Rijn 2019, Evseev and Magor 2021, Rodríguez-Martín D 2021, Wang, Tang et al. 2022, Zheng, Guo et al. 2022). Consequently, our vaccine design should target conserved immunogenic cytotoxic and T-helper cell epitopes to develop the most effective pan-BTV vaccine.

This study predicted highly conserved immunodominant epitopes mainly within NS1, NS2 and NS3 proteins of BTV. These T cell epitopes are considered conserved if they exhibit >60% amino acid conservation (Tengvall 2019, Girdhar, Huang et al. 2022). Our initial analysis confirmed a high degree of conservation, aligning with previous reports (Belaganahalli 2011, Belaganahalli 2012), This finding strongly suggests the existence of conserved CD8+ and CD4+ T cell epitopes within BTV NS proteins. Specifically, we identified these epitopes predominantly in NS1 and NS3. In contrast, NS2 showed variations in epitope presentation between mouse and bovine systems, which is likely influenced by MHC composition and polymorphism. Remarkably, we obtained similar results for CD4+ and CD8+ T cell epitope conservation in both mouse and bovine systems within NS1, NS2, and NS3 proteins. While we could not predict conserved T cell epitopes for the ovine immune system due to current limitations in available tools, our findings suggest that sheep, as natural hosts for BTV, would also recognize and present conserved T cell epitopes to a similar or greater extent as observed in mouse and bovine systems.

In this study, we used the highly conserved epitopes that meet all the criteria for vaccine formulation, including antigenicity, non-allergenicity, and non-toxicity to ensure our vaccine constructs are safe and immunogenic. We paid particular attention to ensure these epitopes are IFNγ inducers, which are known to skew the immune response towards anti-viral immunity (Grandvaux 2002) (E. 2001). Further, we incorporated both CTL and HTL epitopes to boost the effectiveness of the anti-BTV immunity as HTL epitopes are known to induce T helper cells necessary for increasing primary and memory responses of CD8+ CTLs (Umeshappa, Huang et al. 2009{Umeshappa, 2011 #99, Umeshappa 2013, Umeshappa, Xie et al. 2013) .

It is worth noting that all the conserved epitopes identified in NS3 are not inducers of IFNγ but rather stimulate IL-10 responses. This finding aligns with previous experimental reports (Devasthanam 2014) that indicate the suppressive role of the NS3 protein in downregulating IFNγ responses. While LAVs are potent inducers of cellular responses, possibly due to the presence of these proteins, their effectiveness remains limited and serotype specific. Additionally, BTV is known to harbor other negative proteins like VP3 and VP4 (Chauveau, Doceul et al. 2013), which may also impede the development of robust cross-reactive cellular responses against multiple BTV serotypes. These results shed light on why vaccine regimens targeting BTV serotypes have encountered challenges and emphasize the need for meticulous consideration in vaccine design for effective prevention of BTV infections across multiple serotypes. We believe that our thoughtfully formulated vaccine, guided by advanced immunoinformatic tools, holds significant promise for the development of a broad-spectrum vaccine against various BTV serotypes.

Our final vaccine constructs displayed promising attributes in terms of antigenicity, allergenicity, and solubility. Structural modeling results affirmed their acceptable identity and coverage when compared to template structures. The Ramachandran plot analysis reinforced their excellent geometry. Moreover, molecular docking and interaction analyses revealed significant molecular interactions between all four vaccine constructs and TLRs. Consequently, these *in silico*-designed vaccine constructs hold great promise for the development of a broad-spectrum BTV vaccine for bovines. Furthermore, they provide a proof of concept for the development of similar vaccines targeting ovine BTV infections caused by multiple serotypes.

In conclusion, existing BTV vaccines are constrained by serotype-specific limitations, safety concerns, and inadequate cross-protection. Our study explores the potential for a pan-BTV vaccine inspired by the conservation of T cell epitopes in BTV proteins. Computational analysis and *in silico* evaluations of vaccine constructs show promise, although further functional assays and preclinical studies are necessary to validate their efficacy. This approach holds the potential to control the global spread of BTV and prevent outbreaks of novel serotypes, addressing the critical need for broad-spectrum protection against multiple BTV serotypes in ruminants.

## ACKNOWLEDGEMENT

This work is done in collaboration with Centre for Animal Disease Research and Diagnosis, Indian Veterinary Research Institute, Uttar Pradesh, India. The authors thank Ms. Parnian Jahanbani for technical assistance with manuscript formatting. We are grateful for start-up funding by the Department of Microbiology and Immunology, Faculty of Medicine, Dalhousie University. CSU is supported by the prestigious Canada Research Chair Tier 2 Award from Canadian Institutes of Health Research. This work is supported by start-up funding from Dalhousie University. HBK is supported from 2022 DMRF I3V Graduate Studentship. PPCM wishes to acknowledge support from EU H20:20 grant ‘PALE-Blu’ (project number 727393-2).

## AUTHORS CONTRIBUTIONS

H.B.K generated the data in figures 1, 2, 3, 4 and 6, and supplementary table 1-4 with contributions from C.S.U and M.D. M.D generated the data in figure 5 and 7, table 1, and supplementary tables 5-8 with contributions from A.K, and H.B.K under the supervision of D.K and C.S.U. K.P.S, T.K., R.H.N and P.P.C.M contributed to the data interpretation, figures preparation, and manuscript writing. C.S.U. designed the study, supervised, and coordinated its execution, and wrote the manuscript along with H.B.K and A.K.

## SUPPLEMENTARY TABLE LEGENDS

**Supplementary Table 1.** MHC class I specific CD8+ T cell epitopes in the BTV1 proteins.

**Supplementary Table 2.** MHC class II specific CD4+ T cell epitopes in the BTV1 proteins.

**Supplementary Table 3.** BoLA class I specific CD8+ T cell epitopes in the BTV1 proteins.

**Supplementary Table 4.** BoLA class II specific CD4+ T cell epitopes in the BTV1 proteins.

**Supplementary Table 5**. Finalized MHC I and II specific T cell epitopes for the mouse vaccine design.

**Supplementary Table 6**. Finalized BoLA I and II specific T cell epitopes for the bovine vaccine design.

**Supplementary table 7.** Ramachandran plot analysis of the designed vaccine constructs.

