## Supplementary tables for "Immuno-informatics Study Identifies Conserved T Cell Epitopes in Non-structural Proteins of Bluetongue Virus Serotypes: Formulation of Computationally Optimized Next-Generation Broad-spectrum Multiepitope Vaccine"

**Supplementary table 1.** MHC class I specific CD8+ T cell epitopes predicted in the non-structural proteins (NS1, NS2, and NS3) of BTV1.

| **Allele** | **Peptide** | **ic50** | **Percentile** | **% Conservation** |
| --- | --- | --- | --- | --- |
| **NS1** | | | | |
| H-2-Kb | AMPYIYVPI | 185.82 | 0.25 | 88.88 % |
| H-2-Db | ASMAIQQYL | 204.74 | 0.09 | 88.88 % |
| H-2-Kb |  | 494.25 | 0.56 |  |
| H-2-Db | GQIVNPTFI | 19.18 | 0.03 | 88.88 % |
| H-2-Kb | IAYYFYNPD | 90.64 | 0.15 | 44.44 % |
| H-2-Kb | INFLRMDDF | 268.65 | 0.34 | 88.88 % |
| H-2-Db | IQLINFLRM | 69.23 | 0.04 | 88.88 % |
| H-2-Kb |  | 90.62 | 0.15 |  |
| H-2-Kb | KHFNRYASM | 36.95 | 0.07 | 100 % |
| H-2-Kb | MAIQQYLRV | 195.89 | 0.26 | 88.88 % |
| H-2-Kb | RAYATMFEM | 296.74 | 0.37 | 66.66 % |
| H-2-Db | SALVNSERV | 5.96 | 0.01 | 77.77 % |
| H-2-Kb | TFISRYRQI | 312.33 | 0.38 | 88.88 % |
| H-2-Kb | VMFLPIQLI | 150.2 | 0.22 | 88.88 % |
| H-2-Kb | VNPTFISRY | 420.54 | 0.5 | 88.88 % |
| NS3 | | | | |
| H-2-Kb | MSFTEFSSI | 20.15 | 0.04 | 88.88 % |
| H-2-Kb | SWFKSLNPM | 146.52 | 0.22 | 88.88 % |
| H-2-Kb | RAIIRTTLL | 305.28 | 0.37 | 66.66 % |
| H-2-Kb | AAFASYAEA | 331.24 | 0.41 | 100 % |
| H-2-Db | KAMSNTTGA | 408.25 | 0.12 | 100 % |

**Supplementary table 2.** MHC class II specific CD4+ T cell epitopes predicted in the non-structural proteins (NS1, NS2, and NS3) of BTV1.

| **Allele** | **Peptide Sequence** | **IC50** | **Percentile Rank** | **% Conservation** |
| --- | --- | --- | --- | --- |
| **NS1** | | | | |
| H2-IAb | TNCYTGAEALITTAI | 112.08 | 0.22 | 86.66 |
| H2-IAb | TTNCYTGAEALITTA | 123.01 | 0.22 | 80 |
| H2-IAb | YTTNCYTGAEALITT | 162.54 | 0.44 | 80 |
| H2-IAb | NCYTGAEALITTAIH | 179.41 | 0.50 | 86.66 |
| H2-IAb | YYTTNCYTGAEALIT | 255.59 | 0.87 | 80 |
| H2-IAb | KQNFERAMIAATDAE | 265.37 | 0.93 | 86.66 |
| H2-IAb | VKQNFERAMIAATDA | 267.18 | 0.94 | 80 |
| H2-IAb | CVKQNFERAMIAATD | 355.54 | 1.50 | 80 |
| H2-IAb | CYTGAEALITTAIHI | 389.08 | 1.70 | 93.33 |
| H2-IAb | KVHFAGFAAPACESG | 389.87 | 1.70 | 93.33 |
| H2-IAb | TRTFSAISPQWTCSH | 402.37 | 1.80 | 86.66 |
| H2-IAb | ATRTFSAISPQWTCS | 438.48 | 2 | 86.66 |
| H2-IAb | QNFERAMIAATDAEE | 479.93 | 2.20 | 86.66 |
| H2-IAb | LLRWELTKAQRSALL | 490.27 | 2.30 | 86.66 |
| H2-IAb | VHFAGFAAPACESGE | 491.12 | 2.30 | 93.33 |
| H2-IAb | HFAGFAAPACESGEV | 491.74 | 2.30 | 86.66 |
| H2-IAb | FAGFAAPACESGEVI | 519.50 | 2.40 | 86.66 |
| H2-IAb | VLLRWELTKAQRSAL | 521.09 | 2.40 | 86.66 |
| H2-IAb | EKVHFAGFAAPACES | 537.71 | 2.50 | 93.33 |
| H2-IAb | NATRTFSAISPQWTC | 547.76 | 2.50 | 86.66 |
| H2-IAb | AEKVHFAGFAAPACE | 575.95 | 2.60 | 86.66 |
| H2-IAb | MCVKQNFERAMIAAT | 576.94 | 2.70 | 80 |
| H2-IAb | ANATRTFSAISPQWT | 588.10 | 2.70 | 86.66 |
| H2-IAb | RTFSAISPQWTCSHL | 610.96 | 2.80 | 86.66 |
| H2-IAb | RVLLRWELTKAQRSA | 656.22 | 3 | 86.66 |
| H2-IAb | YAEKVHFAGFAAPAC | 667.11 | 3.10 | 86.66 |
| H2-IAb | LRWELTKAQRSALLR | 681.20 | 3.10 | 86.66 |
| H2-IAb | NFERAMIAATDAEEP | 891.04 | 4.20 | 86.66 |
| NS2 | | | | |
| H2-IAb | ERFMSLSSAMPQASG | 167.74 | 0.45 | 100 |
| H2-IAb | FMSLSSAMPQASGGF | 171.72 | 0.48 | 100 |
| H2-IAb | RFMSLSSAMPQASGG | 185.36 | 0.50 | 100 |
| H2-IAb | EWKFEGVSVTPMATR | 217.76 | 0.68 | 100 |
| H2-IAb | DERFMSLSSAMPQAS | 232.12 | 0.73 | 100 |
| H2-IAb | EEWKFEGVSVTPMAT | 275.93 | 0.98 | 100 |
| H2-IAb | WKFEGVSVTPMATRV | 280.83 | 1 | 100 |
| H2-IAb | WEEWKFEGVSVTPMA | 333.17 | 1.40 | 100 |
| H2-IAb | MSLSSAMPQASGGFD | 344.66 | 1.40 | 100 |
| H2-IAb | KFEGVSVTPMATRVQ | 463.16 | 2.10 | 100 |
| H2-IAb | SLSSAMPQASGGFDR | 591.35 | 2.70 | 100 |
| H2-IAb | RWEEWKFEGVSVTPM | 749.45 | 3.40 | 100 |
| H2-IAb | KDERFMSLSSAMPQA | 755.28 | 3.40 | 100 |
| NS3 | | | | |
| H2-IAb | QPPRYAPSAPMPSSM | 119.62 | 0.22 | 86.66 |
| H2-IAb | PRYAPSAPMPSSMPT | 121.07 | 0.22 | 93.33 |
| H2-IAb | PPRYAPSAPMPSSMP | 128.55 | 0.23 | 93.33 |
| H2-IAb | RYAPSAPMPSSMPTV | 140.71 | 0.31 | 93.33 |
| H2-IAb | SQPPRYAPSAPMPSS | 153.53 | 0.38 | 86.66 |
| H2-IAb | ISQPPRYAPSAPMPS | 203.07 | 0.55 | 80 |
| H2-IAb | QKAEKAAFASYAEAF | 407.66 | 1.80 | 100 |
| H2-IAb | KAEKAAFASYAEAFR | 521.36 | 2.40 | 100 |
| H2-IAb | TQKAEKAAFASYAEA | 549.76 | 2.60 | 100 |
| H2-IAb | YAPSAPMPSSMPTVA | 625.10 | 2.90 | 93.33 |
| H2-IAb | QTQKAEKAAFASYAE | 707.08 | 3.30 | 100 |
| H2-IAb | AEKAAFASYAEAFRD | 899.27 | 4.20 | 100 |
| H2-IAb | TQTQKAEKAAFASYA | 980.96 | 4.60 | 100 |

**Supplementary table 3.** BoLA class I specific CD8+ T cell epitopes predicted in the non-structural proteins (NS1, NS2, and NS3) of BTV1.

| **Allele** | **Peptide** | **ic50** | **Percentile Rank** | **% Conservation** |
| --- | --- | --- | --- | --- |
| **NS1** | | | | |
| BoLA-1:02301 | AKAYKLVEL | 236.18 | 0.06 | 55.55 |
| BoLA-1:02301 | AMYDRETVW | 143.0 | 0.04 | 100 |
| BoLA-4:02401 |  | 144.28 | 0.02 |  |
| BoLA-6:01301 |  | 263.81 | 0.67 |  |
| BoLA-6:01301 | AQRSALLRL | 13.36 | 0.05 | 77.77 |
| BoLA-1:02301 |  | 167.77 | 0.04 |  |
| BoLA-6:01302 |  | 274.22 | 0.04 |  |
| BoLA-6:01301 | ASMAIQQYL | 391.5 | 0.87 | 88.88 |
| BoLA-6:01301 | GIFMLGRVL | 303.34 | 0.73 | 88.88 |
| BoLA-6:01301 | GQIVNPTFI | 347.42 | 0.81 | 88.88 |
| BoLA-6:01301 | IQLINFLRM | 315.72 | 0.76 | 88.88 |
| BoLA-1:02301 | IQQYLRVGY | 463.37 | 0.15 | 88.88 |
| BoLA-1:02301 | KHFNRYASM | 298.22 | 0.08 | 100 |
| BoLA-6:01301 | KMKRCGVQL | 7.87 | 0.03 | 77.77 |
| BoLA-6:01302 |  | 190.17 | 0.03 |  |
| BoLA-6:01301 | KQNFERAMI | 25.55 | 0.1 | 88.88 |
| BoLA-1:02301 | MKRCGVQLL | 225.05 | 0.06 | 77.77 |
| BoLA-6:01301 |  | 498.82 | 1.2 |  |
| BoLA-6:01301 | QQYLRVGYA | 310.4 | 0.75 | 77.77 |
| BoLA-1:02301 | RAYATMFEM | 346.88 | 0.09 | 66.66 |
| BoLA-1:02301 | RKHTCQLCY | 394.85 | 0.12 | 88.88 |
| BoLA-1:02301 | RKYNISGDY | 176.92 | 0.04 | 100 |
| BoLA-1:02301 | RMSQMWMEY | 121.99 | 0.03 | 88.88 |
| BoLA-6:01301 | TMFEMVRCI | 435.85 | 0.95 | 77.77 |
| BoLA-1:02301 | VKQNFERAM | 267.1 | 0.07 | 77.77 |
| BoLA-6:01301 | VMFLPIQLI | 209.79 | 0.54 | 88.88 |
| BoLA-6:01301 | WQSWLLPMI | 273.83 | 0.69 | 77.77 |
| NS2 | | | | |
| BoLA-6:01301 | CQIKIGRVI | 119.53 | 0.36 | 88.88 |
| BoLA-1:02301 | IKIGRVIAF | 67.94 | 0.02 | 88.88 |
| BoLA-6:01301 | KLKWQNVPL | 12.07 | 0.05 | 88.88 |
| BoLA-6:01302 |  | 405.49 | 0.05 |  |
| BoLA-6:01301 | KQRKFTKNI | 20.19 | 0.08 | 88.88 |
| BoLA-6:01302 |  | 413.65 | 0.05 |  |
| BoLA-2:01201 | KTLCGVIAK | 307.7 | 0.02 | 88.88 |
| BoLA-1:02301 | LKWQNVPLY | 266.47 | 0.07 | 88.88 |
| BoLA-4:02401 | MIVTKKLKW | 333.96 | 0.02 | 100 |
| BoLA-6:01301 | REMAPRPQV | 476.18 | 1.1 | 55.55 |
| NS3 | | | | |
| BoLA-6:01301 | ALLTSVCTL | 169.04 | 0.47 | 88.88 |
| BoLA-6:01301 | GLKKKRAII | 75.6 | 0.26 | 66.66 |
| BoLA-6:01301 | IQRFEEEKM | 342.28 | 0.8 | 77.77 |
| BoLA-6:01301 | KLKSDLSGL | 79.69 | 0.27 | 55.55 |
| BoLA-6:01301 | KMKHNQDRV | 314.3 | 0.76 | 77.77 |
| BoLA-6:01301 | RAIIRTTLL | 23.82 | 0.1 | 66.66 |
| BoLA-6:01301 | RLRQIKRHV | 202.96 | 0.53 | 100 |

**Supplementary table 4.** BoLA class II specific CD4+ T cell epitopes predicted in the non-structural proteins (NS1, NS2, and NS3) of BTV1.

| **Allele** | **Peptide** | **Core_Rel** | **Score_EL** | **%Rank_EL** | **% Conservation** |
| --- | --- | --- | --- | --- | --- |
| **NS1** | | | | | |
| BoLA-DRB3_2002 | CESGEVINLAARMSQ | 1.000 | 0.839459 | 0.59 | 93.33 |
| BoLA-DRB3_1501 | CIITLCYAEKVHFAG | 1.000 | 0.801402 | 0.46 | 93.33 |
| BoLA-DRB3_0101 | EALITTAIHIHRWIR | 0.987 | 0.798823 | 0.48 | 93.33 |
| BoLA-DRB3_1001 | FAKHFNRYASMAIQQ | 1.000 | 0.552939 | 0.77 | 93.33 |
| BoLA-DRB3_1601 | FSQPYEEDVEGKMKR | 1.000 | 0.711918 | 0.65 | 86.66 |
| BoLA-DRB3_1201 | HFNRYASMAIQQYLR | 0.967 | 0.672970 | 0.92 | 93.33 |
| BoLA-DRB3_1501 | KQSALVNSERVRLDD | 1.000 | 0.958137 | 0.10 | 86.66 |
| BoLA-DRB3_1501 | LRWELTKAQRSALLR | 1.000 | 0.708936 | 0.75 | 86.66 |
| BoLA-DRB3_1201 | PYIYVPIKEGQIVNP | 0.987 | 0.669020 | 0.94 | 93.33 |
| BoLA-DRB3_1601 |  | 0.993 | 0.894352 | 0.19 |  |
| BoLA-DRB3_1101 | QSALVNSERVRLDDS | 0.980 | 0.573046 | 0.86 | 86.66 |
| BoLA-DRB3_1101 | QSWLLPMIIVREGLD | 0.953 | 0.630295 | 0.69 | 73.33 |
| BoLA-DRB3_1201 | RQIAYYFYNPDAADD | 0.900 | 0.751418 | 0.65 | 66.66 |
| BoLA-DRB3_2002 | SAMPYIYVPIKEGQI | 0.940 | 0.887746 | 0.38 | 86.66 |
| BoLA-DRB3_1001 | SMAIQQYLRVGYAEE | 1.000 | 0.501013 | 0.93 | 86.66 |
| NS2 | | | | | |
| BoLA-DRB3_1601 | FTKNIFVLDVNAKTL | 1.000 | 0.922736 | 0.14 | 80 |
| BoLA-DRB3_1001 | GFDRMIVTKKLKWQN | 0.993 | 0.845177 | 0.18 | 93.33 |
| BoLA-DRB3_1501 |  | 0.993 | 0.781267 | 0.52 |  |
| BoLA-DRB3_1601 | HDDFRADEDERGEDA | 1.000 | 0.695474 | 0.71 | 26.66 |
| BoLA-DRB3_0101 | KEYIEKVAKQVKLKD | 0.993 | 0.987052 | 0.01 | 80 |
| BoLA-DRB3_1001 | MGIVQPYMKNDFDRN | 1.000 | 0.657376 | 0.51 | 93.33 |
| BoLA-DRB3_1501 | PYCQIKIGRVIAFKP | 0.993 | 0.778683 | 0.53 | 86.66 |
| BoLA-DRB3_1501 | RLEGLREARANIFKE | 1.000 | 0.787897 | 0.50 | 60 |
| NS3 | | | | | |
| BoLA-DRB3_1201 | FTEFSSIPLDGFEMP | 1.000 | 0.664043 | 0.95 | 80 |
| BoLA-DRB3_1201 | LRQIKRHVNEQILPK | 1.000 | 0.701043 | 0.83 | 100 |
| BoLA-DRB3_1001 | NQQIDMIKKEVMKKQ | 1.000 | 0.512959 | 0.89 | 86.66 |
| BoLA-DRB3_1501 |  | 1.000 | 0.923153 | 0.17 |  |
| BoLA-DRB3_1201 | SGLIQRFEEEKMKHN | 1.000 | 0.897994 | 0.26 | 86.66 |
| BoLA-DRB3_1501 | VNEQILPKLKSDLSG | 1.000 | 0.765585 | 0.57 | 73.33 |
| BoLA-DRB3_1001 | VRMSFTEFSSIPLDG | 1.000 | 0.833452 | 0.20 | 93.33 |
| BoLA-DRB3_2002 | PSWFKSLNPMLGVVN | 1.000 | 0.922687 | 0.25 | 80 |
| BoLA-DRB3_2002 | LGATFLMMVCAKSER | 1.000 | 0.826423 | 0.64 | 80 |

**Supplementary table 5.** Finalized MHC I and II specific T cell epitopes for the mouse vaccine design.

| **CD8+ T cell epitopes** | **NS1** | | | | | | |
| --- | --- | --- | --- | --- | --- | --- | --- |
|  | **Allele** | **Epitope** | **Antigenicity score** | **Antigenicity** | **Allergenicity** | **Toxicity** | |
|  | **H-2-Kb** | INFLRMDDF | 1.0767 | Antigen | NON-Allergen | Non-Toxin | |
|  | **H-2-Db**  **H-2-Kb** | IQLINFLRM | 0.744 | Antigen | NON-Allergen | Non-Toxin | |
|  | **H-2-Kb** | TFISRYRQI | 0.6903 | Antigen | NON-Allergen | Non-Toxin | |
|  | **H-2-Kb** | VMFLPIQLI | 1.0511 | Antigen | NON-Allergen | Non-Toxin | |
|  | **H-2-Kb** | VNPTFISRY | 1.7189 | Antigen | NON-Allergen | Non-Toxin | |
| **NS1** | | | | | | | |
| **CD4+ T cell epitopes** | **Allele** | **Epitope** | **Antigenicity score** | **Antigenicity** | **Allergenicity** | **Toxicity** | **IFNg** |
|  | **H2-IAb** | ATRTFSAISPQWTCS | 0.9288 | Antigen | NON-Allergen | Non-Toxin | POSITIVE |
|  | **H2-IAb** | NATRTFSAISPQWTC | 1.0800 | Antigen | NON-Allergen | Non-Toxin | POSITIVE |
|  | **Allele** | **Epitope** | **Antigenicity score** | **Antigenicity** | **Allergenicity** | **Toxicity** | **IFNg** |
|  | **H2-IAb** | EWKFEGVSVTPMATR | 1.5224 | Antigen | NON-Allergen | Non-Toxin | POSITIVE |
|  | **H2-IAb** | WKFEGVSVTPMATRV | 1.2000 | Antigen | NON-Allergen | Non-Toxin | POSITIVE |
|  | **H2-IAb** | KFEGVSVTPMATRVQ | 1.0904 | Antigen | NON-Allergen | Non-Toxin | POSITIVE |

**Supplementary table 6.** Finalized BoLA I and II specific T cell epitopes for the bovine vaccine design.

| **NS1** | | | | | | | |
| --- | --- | --- | --- | --- | --- | --- | --- |
| **CD8+ T cell epitopes** | **Allele** | **Epitope** | **Antigenicity score** | **Antigenicity** | **Allergenicity** | **Toxicity** | |
|  | BoLA-6:01301 | IQLINFLRM | 0.7444 | Antigen | Non-allergen | Non-Toxin | |
|  | BoLA-6:01301 | QQYLRVGYA | 1.2748 | Antigen | Non-allergen | Non-Toxin | |
|  | BoLA-1:02301 | RKYNISGDY | 0.6154 | Antigen | Non-allergen | Non-Toxin | |
|  | BoLA-6:01301 | VMFLPIQLI | 1.0511 | Antigen | Non-allergen | Non-Toxin | |
|  | **NS2** | | | | | | |
|  | BoLA-1:02301 | IKIGRVIAF | 1.0138 | Antigen | Non-allergen | Non-Toxin | |
|  | BoLA-6:01301 | KLKW**Q**NVPL | 2.2848 | Antigen | Non-allergen | Non-Toxin | |
|  | BoLA-1:02301 | LKW**Q**NVPLY | 1.7890 | Antigen | Non-allergen | Non-Toxin | |
|  | BoLA-4:02401 | MIVTKKLKW | 1.7636 | Antigen | Non-allergen | Non-Toxin | |
| **NS1** | | | | | | | |
| **CD4+ T cell epitopes** | **Allele** | **Epitope** | **Antigenicity score** | **Antigenicity** | **Allergenicity** | **Toxicity** | **IFNg** |
|  | BoLA-DRB3_1501 | CIITLCYAEKVHFAG | 0.5405 | Antigen | Non-allergen | Non-Toxin | Positive |
|  | BoLA-DRB3_1501 | KQSALVNSERVRLDD | 0.7360 | Antigen | Non-allergen | Non-Toxin | Positive |
|  | BoLA-DRB3_1201 | PYIYVPIKEGQIVNP | 0.7904 | Antigen | Non-allergen | Non-Toxin | Positive |
|  | BoLA-DRB3_1101 | QSWLLPMIIVREGLD | 0.5591 | Antigen | Non-allergen | Non-Toxin | Positive |
|  | BoLA-DRB3_1001 | SMAIQQYLRVGYAEE | 0.8971 | Antigen | Non-allergen | Non-Toxin | Positive |

**Supplementary table 7.** Ramachandran plot analysis of the designed vaccine constructs.

|  | **Mouse vaccine with TLR3 adjuvant** | **Mouse vaccine with TLR4 adjuvant** | **Bovine vaccine with TLR3 adjuvant** | **Bovine vaccine with TLR3 adjuvant** |
| --- | --- | --- | --- | --- |
| **Favoured region** | 86.9 % | 92.7 % | 90.8 % | 96 % |
| **Additionally allowed region** | 10.4 % | 6.1 % | 7.2 % | 3 % |
| **Generally allowed region** | 0.5 % | 0 % | 1.4 % | 0 % |
| **Disallowed** | 0 % | 1.2 % | 0.5 % | 1 % |
